# Laser capture microdissection coupled mass spectrometry (LCM-MS) for spatially resolved analysis of formalin-fixed and stained human lung tissues

**DOI:** 10.1101/721373

**Authors:** Jeremy A. Herrera, Venkatesh Mallikarjun, Silvia Rosini, Maria Angeles Montero, Stacey Warwood, Ronan O’Caulian, David Knight, Martin A. Schwartz, Joe Swift

**Affiliations:** The Wellcome Centre for Cell-Matrix Research, University of Manchester, Manchester, M13 9PT, UK; Division of Cell Matrix Biology and Regenerative Medicine, School of Biological Sciences, Faculty of Biology, Medicine and Health, University of Manchester, Manchester Academic Health Science Centre, Manchester, M13 9PL, UK; Histopathology Department, Manchester University NHS Foundation Trust, Southmoor Road, Wythenshawe, Manchester, M23 9LT, UK

## Abstract

Haematoxylin and eosin (H&E) – which respectively stain nuclei blue and other cellular and stromal material pink – are routinely used for clinical diagnosis based on the identification of morphological features. A richer characterization can be achieved by laser capture microdissection coupled to mass spectrometry (LCM-MS), giving an unbiased assay of the proteins that make up the tissue. However, the process of fixing, and H&E staining of tissues is poorly compatible with standard sample preparation methods for mass spectrometry, resulting in low protein yield. Here we describe a microproteomics technique optimized to analyze H&E-stained, formalin-fixed paraffin-embedded (FFPE) tissues. We advance our methodology by combining 3 techniques shown to individually enhance protein yields (heat extraction, physical disruption, and in column digestion) into one optimized pipeline for the analysis of H&E stained FFPE tissues. Micro-dissected morphologically normal human lung alveoli (0.082 mm^3^) and human lung blood vessels (0.094 mm^3^) from FFPE fixed section from Idiopathic Pulmonary Fibrosis (IPF) specimens were then subject to comparative proteomics using this methodology. This approach yielded 1252 differentially expressed proteins including 137 extracellular matrix (ECM) proteins. In addition, we offer proof of principal that MS can identify distinct, characteristic proteomic compositions of anatomical features within complex tissues.

## INTRODUCTION

Mass spectrometry (MS) proteomics is a powerful tool to systemically identify and quantify proteins in complex biological samples. The utility of this method is maximized when performed with spatial resolution to report on the composition and function of specific regions of tissue. Extracellular matrix (ECM) is particularly important in determining cell behaviour in health and disease (1) but is especially challenging for proteomic analysis given the extensive covalent crosslinking and low solubility of many ECM proteins (2). However, common protocols for bottom-up proteomics (i.e. based on detection of peptide protein fragments) require sample homogenization and digestion, resulting in a loss of any information regarding protein localization and spatial relationships. To this end, laser capture microdissection coupled to mass spectrometry (LCM-MS) is a method currently being optimized for microproteomics to determine regional tissue differences (3). LCM-MS has been performed using fresh (4), flash-frozen (5–7), and formalin-fixed paraffin-embedded (FFPE) tissues (8–11). In this study, we describe and examine the performance of a protocol for LCM-MS analysis of FFPE sections of human lung tissue that were haematoxylin and eosin (H&E) stained.

H&E staining provides critical morphological characterization enabling researchers to identify anatomical features of interest. However, haematoxylin staining has been shown to reduce protein detection by MS (12). Currently, few LCM-MS studies exist for H&E stained FFPE tissue sections. In an earlier study, up to 866 proteins (13) were identified from H&E stained FFPE sections of human head and neck squamous cell carcinomas. In a recent study, using 0.5 mm^3^ of tissue, researchers identified up to 714 unique proteins from H&E stained FFPE tissue sections of cutaneous squamous cell carcinoma (14). The need for novel LCM-MS protocols for H&E stained FFPE tissue is warranted.

Herein, we describe and demonstrate the application of a protocol for microproteomics that combines multiple steps that individually enhance protein yield to maximize protein retrieval from H&E-stained, FFPE tissue. First, we perform a detergent-based heat-retrieval procedure that was shown to enhance protein solubility (15–17). We combined this with two techniques to enhance extraction of ECM proteins: physical disruption (10, 18) and chemical extraction with a urea-based buffer (19). Lastly, we utilize an in-column trypsin-digest system [a recently commercialized product, S-Trap (20)], to increase peptide yields (12) while effectively removing detergents and contaminants from the samples (21). These combined methods enabled us to identify 137 ECM proteins and 1252 proteins that were differentially expressed (with a threshold of three or more unique identifying peptides) between samples of morphologically normal human lung alveoli (0.082 mm^3^ of tissue used) and human lung blood vessels (0.094 mm^3^ of tissue used) from formalin-fixed, paraffin-embedded and H&E stained sections. Data analysis demonstrated enrichment of cellular and ECM protein components characteristic of the different regions of tissue.

## MATERIALS AND METHODS

### Procurement of human lung tissue

The use of human lung tissue was approved by University of Manchester Health Research Authority with patient consent under protocol REC#14/NW/0260. The specimens used for this study met the criteria for Idiopathic Pulmonary Fibrosis (IPF) diagnosis (22), however, we used distal lung tissue that appeared morphologically normal for the LCM-MS study.

### Immunohistochemistry

Human lung samples were formalin-fixed and paraffin-embedded (FFPE). Deparaffinized and rehydrated 5-micron sections were subjected to antigen heat retrieval using citrate buffer (Abcam, ab208572), for 30 minutes at 100 °C, cooled to room temperature for 20 minutes, treated with 3% hydrogen peroxide for 5 minutes, blocked in TBS Super Block for 1 hour (Thermo Fisher; 37581), and probed with primary antibody (TNC, 1:500, Abcam, ab108930; LAMB1, 1:1,000, Abcam, ab16048) overnight in 10% blocking solution. The following day, the specimens were subjected to Novolink Polymer Detection Systems (Leica RE7270-RE, per the manufacturer’s recommendations), developed for 5 minutes with DAB Chromagen (Cell Signal, 11724), counterstained with haematoxylin and cover-slipped with Permount (Thermo Fisher Scientific, SP15).

### Pentachrome staining

We followed a modified Russell-Movats pentachrome staining protocol. Deparaffinized specimens were stained with alcian blue for 20 minutes (1% alcian blue [Sigma-Aldrich, A-1986] and 1% glacial acetic acid), treated in alkaline alcohol for 1 hour (90% alcohol and 10% of a 30% ammonium hydroxide solution [Sigma-Aldrich, 221228]), alcohol haematoxylin solution for 10 minutes (50% of a 5% absolute alcoholic haematoxylin [Sigma Aldrich, P4006], 25% of a 10% aqueou ferric chloride [Sigma-Aldrich, F2877], and 25% of 2 grams Iodine [Alfa Aesar, A12278], 4 grams potassium iodide [Fluorochem, 319032] in 100 mL water), 5% aqueous sodium thiosulfate for 1 minute (Sigma-Aldrich, S-6672), and then crocein scarlet-acid fuchsin solution for 2 minutes (4 parts crocein scarlet – 0.1% crocein scarlet [Alfa Aesar, J66876] and 0.5% glacial acetic acid; 1 part acid fuchsin – 0.1% acid fuchsin [Alfa Aesar, B22222] and 0.5% glacial acetic acid). Sections were then treated 2 times with 5% phosphotungstic acid for 5 minutes each (Sigma Aldrich, P4006), washed 3 times with 100% alcohol, stained with 6% alcoholic saffron (VWR, 283-295-0) for 15 minutes, and then coverslipped with Permount.

### Haematoxylin and eosin staining

5-micron FFPE sections were mounted onto MMI membrane slides (MMI, 50102) and stained using an automated stainer (Leica XL) at the Histology Core at University of Manchester. In short, the FFPE slides were dewaxed by xylene and alcohol treatment, followed by a 2-minute hematoxylin incubation, acid alcohol treated, and stained with eosin for 1 minute. Slides were then washed in 100% ethanol and allowed to air dry. Slides were then stored in 4 °C for up to one week before LCM-MS.

### Histological Imaging

Stained slides were imaged using a DMC2900 Leica camera along with Lecia Application Suite X software (Leica).

### Laser capture microdissection

The 5-micron H&E sections were loaded onto the MMI CellCut Laser Microdissection system (Molecular Machines & Industries). We used a power setting of 75% laser moving at 40 microns/second for automated cutting and collected laser microdissected specimens using MMI transparent caps (MMI, 50204).

### Sample preparation for mass spectrometry

Laser microdissected tissue was resuspended in 25 µL 50 mM triethylammonium bicarbonate (TEAB) (Sigma, T7408), 5% SDS (pH 7.5) and subjected to 95°C for 20 minutes, then 60 °C for 2 hours while shaking at 1400 RPM (Eppendorf, ThermoMix C). To select for ECM proteins, we then added 75 µL of a 50 mM TEAB, 5% SDS, 10 M urea (pH 7.5) solution to the 25 µL sample, after it had cooled to room temperature, to create a final volume of 100 µL of 50 mM TEAB, 5% SDS, 7.5 M urea (pH 7.5). Samples were then placed into a Covaris microtube (Covaris, 520045) and sheared using the LE220-Plus Focused Ultrasonicator (Covaris, UK) set at 6 °C with the following settings: duration of 50 seconds, peak power 500, duty factor of 20.0%, cycles/burst of 200, average power at 100, and then delayed for 10 seconds. This was repeated for a total of 10 cycles (10-minute total run time). After shearing, samples were alkylated by the addition of 8 µL of 500 mM iodoacetamide (Sigma, I1149) and incubated for 30 minutes in the dark. Samples were then acidified by the addition of 12 µL of 12% aqueous phosphoric acid (Sigma, 345245) and centrifuged at 12000 RPM for 5 minutes. The supernatant was collected and resuspended with 600 µL of 90% methanol, 100 mM TEAB (pH 7.10). The sample was then added to a S-Trap column (ProtiFi, C02-micro) and centrifuged at 2000 RPM using 200 µL at a time until all the sample had passed through the column. After discarding the flow through, the S-Trap column was washed by adding 150 µL of 90% methanol, 100 mM TEAB (pH 7.10) and centrifuging at 2000 RPM. Washing was repeated a further 9 times, discarding the flow through each time. In-column digest was performed by adding 25 µL of a solution containing 2 µg trypsin (Promega, V5111) in 50 mM TEAB pH 8.0. Trypsin digestion was performed at 47 °C for 1 hour without shaking. Samples were eluted by adding 40 µL of 50 mM TEAB (pH 8.0) and centrifuging at 2000 RPM, followed by the addition of 40 µL of 0.2% aqueous formic acid (Sigma Aldrich, 27001) and centrifuging, and finally adding 50% aqueous acetonitrile (Fisher, A955-212) and centrifuging. Eluted fractions were combined and the total 120 µL sample was then lyophilised using a speed-vac (Heto Cooling System).

Desalting of samples was performed using Oligo R3 resin beads. Briefly, 100 µL of a 10 mg/mL Oligo R3 resin (Thermo Scientific, 1-1339-03) in aqueous 50% acetonitrile was placed into a 96-well 0.2 µm PVDF filter plate (Corning, 3504). The plate was centrifuged at 1400 RPM (Thermo Scientific, Megafuge 16) for 1 minute to clear Oligo R3 resin with a blank 96-well plate underneath to catch the flow through which was then discarded. 100 µL of aqueous 50% acetonitrile was mixed with the resin and centrifuged again, discarding the flow through. Finally, 100 µL of aqueous 0.1% formic acid were mixed with the resin and centrifuged for a total of two repeats, while discarding the flow through. Samples were then resuspended in 100 µl of aqueous 5% acetonitrile, 0.1% formic and mixed with the now washed Oligo R3 Resin and allowed to shake on a plate shaker (Eppendorf, Thermomixer Comfort) for 5 minutes at 800 RPM, and then centrifuged (flow through was discarded). The sample peptides were now bound to the Oligo R3 Resin and washed for a total of ten times by the addition of 100 µL of aqueous 0.1% formic acid, mixed for 2 minutes at 800 RPM, centrifuged, and flow through discarded. Finally, the washed peptides were eluted by mixing with 50 µL of aqueous 50% acetonitrile for 2 minutes, centrifuged, collecting the flow through in a clean 96-well capture plate. Elution was repeated with an additional 50 µL of aqueous 50% acetonitrile and retained. Desalted peptides were lyophilised in a speed-vac and stored at 4 °C until needed.

### Liquid chromatography coupled tandem mass spectrometry

Peptides were evaluated by liquid chromatography (LC) coupled tandem MS (LC-MS/MS) using an UltiMate 3000 Rapid Separation LC system (RSLC, Dionex Corporation) coupled to a Q Exactive HF (Thermo Fisher). Peptide mixtures were separated with a multistep gradient from 95% (0.1% aqueous formic acid, FA) and 5% A (0.1% FA in acetonitrile), to 7% A at 1 min, 18% A at 58 min, 27% A at 72 min and 60% A at 74 min at 300 nL/min, using a 75 mm × 250 μm, inner diameter = 1.7 μm, CSH C18, analytical column (Waters). Peptide fragmentation was performed automatically by data dependent analysis and chromatograms were created using Xcalibur software.

### Mass spectrometry data analysis and statistics

Raw spectra were automatically aligned using Progenesis QI for proteomics (version 4.1; Nonlinear Dynamics, Waters). Progenesis QI’s alignment feature allows MS2 spectral information to be shared across samples so that MS1 spectra that do not have associated MS2 spectra in a given sample (owing to low abundance and thus not being selected for fragmentation in data-dependent acquisition) can still be identified. Alignment decreases the number of missing values (a common problem in proteomics) at the risk of wrongly inferring presence of a peptide when it may genuinely be absent. Spectra from different tissue sections (human lung blood vessel or morphologically normal human lung alveoli) were analyzed either separately (with alignment between donors but not between sections; **Fig. 2**) or together (with alignment between both sections and donors; **Fig. 3**). Peak-picking in Progenesis QI was performed using default parameters and subsequent features were filtered to leave only peptides with a charge of +1 to +4, with 3 or more isotopes. Remaining features were searched using Mascot (server version 2.5.1, parser version 2.5.2.0; Matrix Science), against the SwissProt and TREMBL human database. The peptide database was modified to search for alkylated cysteine residues (monoisotopic mass change, +57.021 Da) as a fixed modification, with oxidized methionine (+15.995 Da), hydroxylation of asparagine, aspartic acid, proline or lysine (+15.995 Da) and phosphorylation of serine, tyrosine, threonine (+79.966 Da) as variable modifications. A maximum of two missed cleavages was allowed. Peptide tolerance and MS/MS tolerance were set to 8 ppm and 0.015 Da, respectively. Peptide detection intensities were exported from Progenesis QI as comma separated variable (.csv) spreadsheets for further processing. Peptides assigned to proteins with ‘unreviewed’ status in the UniProt database were reassigned to the most abundant ‘reviewed’ protein with sequence identity in the dataset. Peptides shared between different protein groups were excluded from subsequent analysis. For differential expression analysis in **Fig. 3**, fold changes were calculated using the Matlab (version 2015a; The MathWorks) implementation of BayesENproteomics (23), available from: https://github.com/VenkMallikarjun/BayesENproteomics. BayesENproteomics fits regularized regression models to take into account donor variability and variable behaviour of peptides assigned to a single protein group (due to post-translational modifications or differential splicing), weighting observations based on confidence in peptide identification (inferred via Mascot scores). These features allow BayesENproteomics to calculate fold changes for the dominant proteoforms of proteins represented in a complex clinical dataset. Missing values were imputed within BayesENproteomics model fitting using an adaptive multiple imputation method that attempts to discern whether a given missing value is missing at random (MAR) or non-randomly (MNR) and imputes from appropriate distributions (23). Reactome (24, 25) Pathway enrichment analysis was performed as described in (23), by fitting linear models for each pathway represented in the dataset, based on protein-level fold changes calculated as described above.

**Figure 1:**
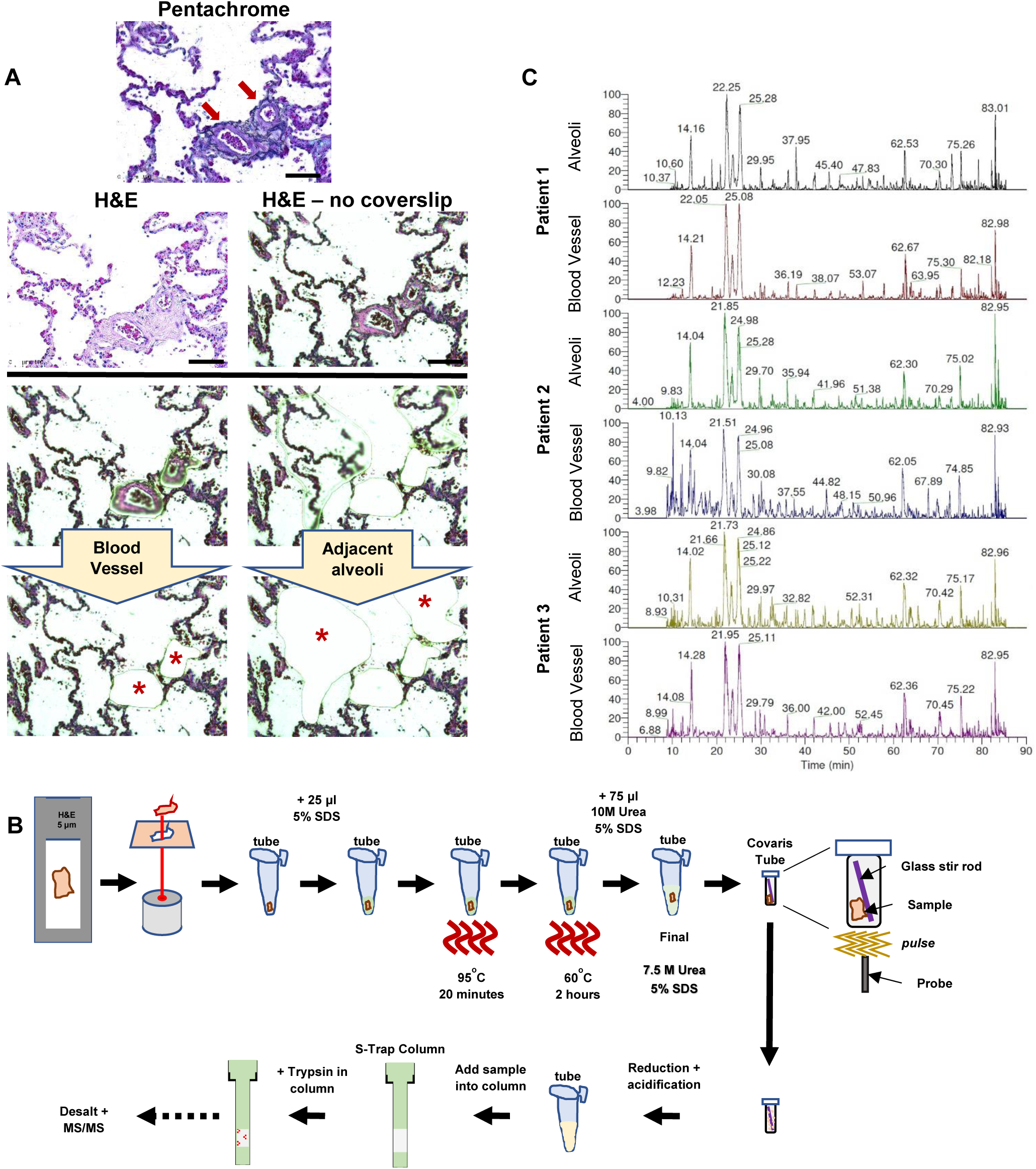
Laser capture microscopy of human lung blood vessels for mass spectrometry. (**A**) Blood vessels are identified by pentachrome stain (red arrows) and serial sections are used to laser microdissect adjacent morphologically normal human lung alveoli and human lung blood vessels. Scale bar represents 100 μm. (**B**) A workflow of tissue preparation for mass spectrometry. Laser captured tissue is detergent treated, subjected to heat, resuspended with urea, sheared using Covaris instrument, and samples are later placed into a S-Trap Column for trypsin digest, followed by desalting prior to mass spectrometry loading. (**C**) Chromatograms are shown for morphologically normal human lung alveoli and human lung blood vessels for each patient (n = 3).

**Figure 2:**
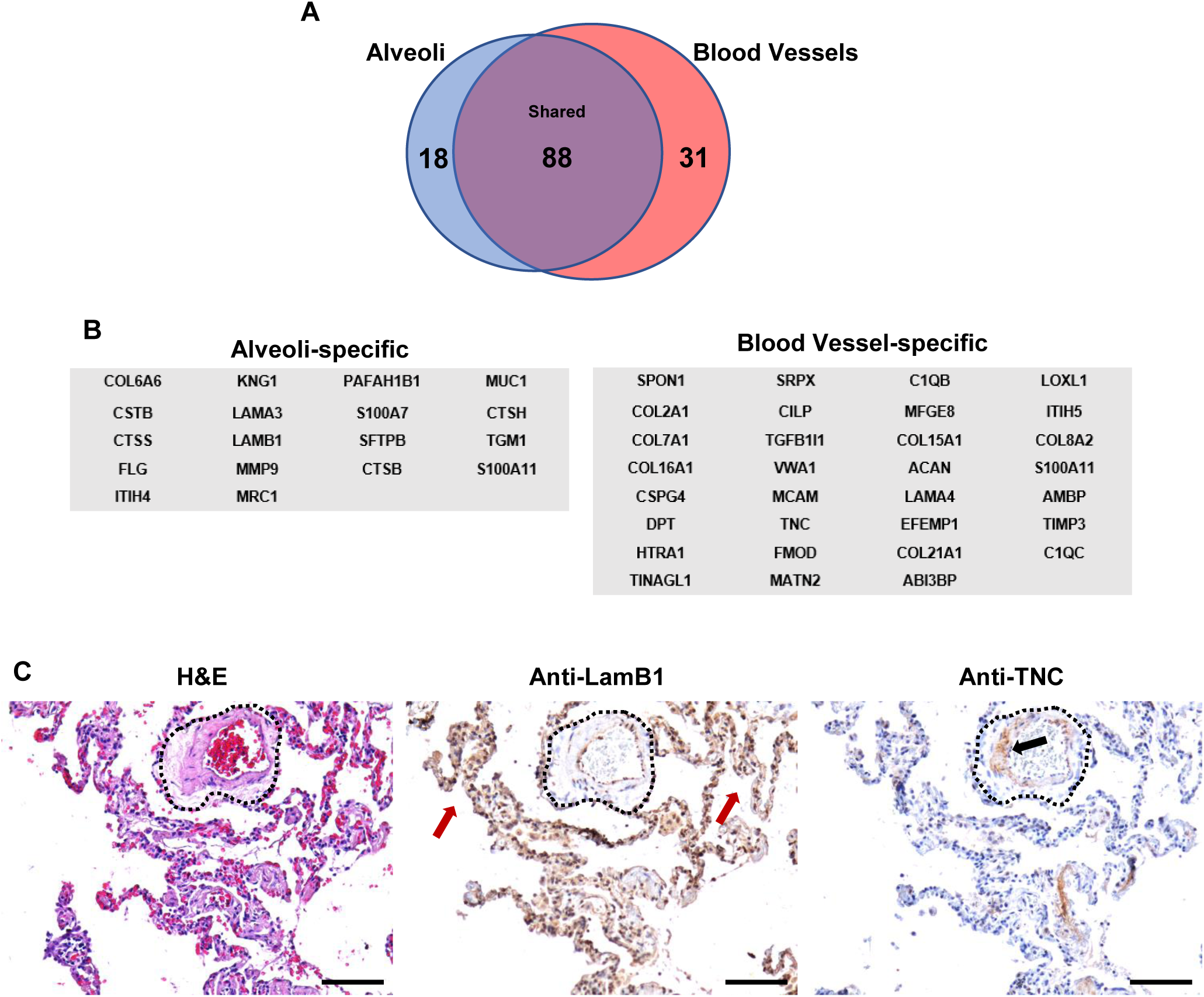
The ECM comprising morphologically normal alveoli and blood vessels in IPF. (**A**) A Venn diagram showing the number of ECM constituents within morphologically normal human lung alveoli and human lung blood vessels (n = 3 IPF specimens). (**B**) A list of ECM genes specific for morphologically normal alveoli and blood vessels. (**C**) Serial sections stained with H&E, anti-LamB1 and anti-TNC. A blood vessel is outlined in black dots, red arrows highlight intense immunostain for LamB1, and black arrow highlights immunostain for TNC within the blood vessel. Scale bar represents 100 μm.

**Figure 3:**
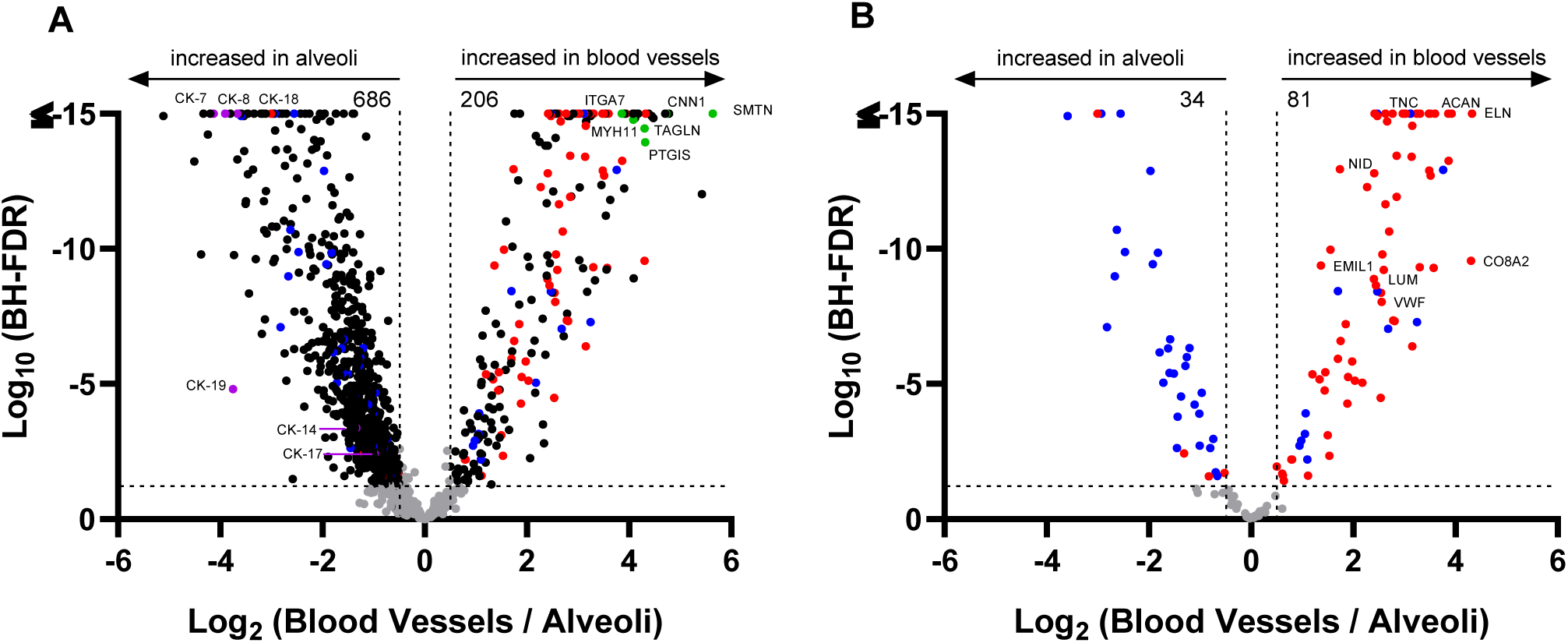
The ECM comprising morphologically normal human lung alveoli and human lung blood vessels in IPF. (**A**) A volcano plot of all 1252 proteins or (**B**) ECM proteins only showing a negative natural log of the FDR values plotted against the base 2 log of the change for each protein. The thresholds are set for a base log 2 >0.5 and FDR p-value < 0.05; n = 3 IPF specimens. Purple dots indicate known proteins expressed in alveoli, green dots indicate known proteins expressed in blood vessels, blue dots represent matrisome-associated proteins, and red dots represent core-matrisome proteins.

### Data availability

Raw mass spectrometry data were deposited to ProteomeXchange with the identifier PXD014762.

## RESULTS

### Optimizing microproteomics for haematoxylin & eosin-stained formalin-fixed paraffin-embedded tissue

Morphologically normal human lung alveoli and human lung blood vessels were laser microdissected from uninvolved IPF tissue (**Fig 1A**) using a pentachrome and H&E stain as a guide. Per specimen (n = 3 IPF patients), approximately 0.082 mm^3^ of morphologically normal human lung alveoli and 0.094 mm^3^ of human lung blood vessels were pooled from H&E-stained 5-micron FFPE sections and used for downstream mass spectrometry; encompassing a total of 6 LCM-MS samples (Schematic in **Fig 1B**). In short, samples were resuspended in 5% SDS, and heated to denature proteins, then resuspended at room temperature in a final solution of 7.5 M urea and 5% SDS to enhance ECM solubility (19). Samples were then sheared using a LE220 focused-ultrasonicator (Covaris Ltd, United Kingdom). This combined protocol extracted more proteins as assessed by SDS-PAGE than individual methods (**Supplemental Figure 1**). Samples were then placed into an S-Trap column (ProtiFi, NY, USA) (20), desalted, and analyzed for mass spectrometry. To show that similar peptide loading was achieved, we show the chromatograms of the 6 samples (3 IPF specimens stratified into morphologically normal human lung alveoli and human lung blood vessels) (**Fig 1C**).

### The ECM comprising morphologically normal human lung alveoli and blood vessels

We next used Progenesis QI (Nonlinear Dynamics) and Mascot (Matrix Science) to process the data acquired from the MS analysis. With a cut-off of 3 or more peptides, we identified 1107 and 683 proteins in morphologically normal human lung alveoli and human lung blood vessels, respectively. We then used the Human Matrisome Project (http://matrisomeproject.mit.edu) which provides a list of core matrisome proteins (ECM glycoproteins, collagens, and proteoglycans) and matrisome-associated proteins (ECM-affiliated proteins, ECM regulators, and secreted factors) and directly compared to our protein list (26). This analysis identified 106 and 119 ECM proteins in morphologically normal human lung alveoli and human lung blood vessels, respectively. We show that 88 ECM constituents are shared within these tissues (**Supplemental Table 1**), with 18 proteins unique to morphologically normal human lung alveoli and 31 proteins unique to human lung blood vessels (**Fig 2A**). The protein gene names for these regions are shown in **Fig 2B**. To validate these results, we carried out immunostaining for two proteins that proteomics identified as specific to each tissue. Antibody to laminin subunit beta-1 (LAMB1) stained predominantly the alveoli (red arrows), whereas antibody to tenascin (TNC) stained blood vessels but not alveoli (blood vessel outlined in black dots; **Fig 2C**).

To gain further insight into the differences in the total protein abundance between morphologically normal human lung alveoli and human lung blood vessel, we used Progenesis QI to align all samples from different sections together, followed by BayesENproteomics (23) to compare the differential expression of the 1252 proteins identified (at a protein identification threshold of 3 unique peptides). This analysis showed that 206 proteins are enriched in blood vessels whereas 686 are enriched in morphologically normal alveoli (**Fig 3A**). In accordance with prior knowledge, proteins known to be expressed in blood vessels, such as smoothelin, CNN1, ITGA7, MYH11, TAGLN, and PTGIS are over-represented in the blood vessels (highlighted in green dots) (27–30). The top 10 pathways and top 10 proteins enriched in human lung blood vessels are shown in **Tables 1 & 2**, respectively. Similarly, cytokeratin-7, −8, −18, −19, 14, and −17 are enriched in the alveoli as previously shown (highlighted in purple dots) (31, 32). The top 10 pathways and top 10 proteins enriched in alveoli are shown in **Tables 3 & 4**, respectively. These data suggest that our methodology of laser capture microscopy coupled with mass spectrometry can robustly discriminate regional proteomics in the same specimen.

**Table 1:** Reaction pathways enriched in blood vessels.

| **Pathway Name/**<br>**Reactome Pathway Identifier (R-HSA)** | **Log2 Fold-**<br>**Change** | **FDR** | **Set Size** |
| --- | --- | --- | --- |
| Collagen chain trimerization/<br>R-HSA-8948216 | 2.71 | 1.83E-14 | 22 |
| ECM proteoglycans/<br>R-HSA-3000178 | 2.20 | 2.12E-14 | 36 |
| Collagen biosynthesis and modifying<br>enzymes/ R-HSA-1650814 | 2.23 | 2.22E-11 | 26 |
| Non-integrin membrane-ECM interactions/<br>R-HSA-300171 | 2.36 | 9.69E-11 | 20 |
| Molecules associated with elastic fibres/<br>R-HSA-2129379 | 2.21 | 1.11E-10 | 15 |
| Syndecan interactions/<br>R-HSA-3000170 | 2.45 | 1.11E-10 | 12 |
| Extracellular matrix organization/<br>R-HSA-1474244 | 2.84 | 3.42E-10 | 10 |
| MET activates PTK2 signaling/<br>R-HSA-8874081 | 2.26 | 9.02E-10 | 15 |
| Collagen degradation/<br>R-HSA-1442490 | 2.18 | 3.12E-09 | 16 |
| Laminin interactions/<br>R-HSA-3000157 | 1.95 | 2.36E-08 | 6 |

**Table 2:**
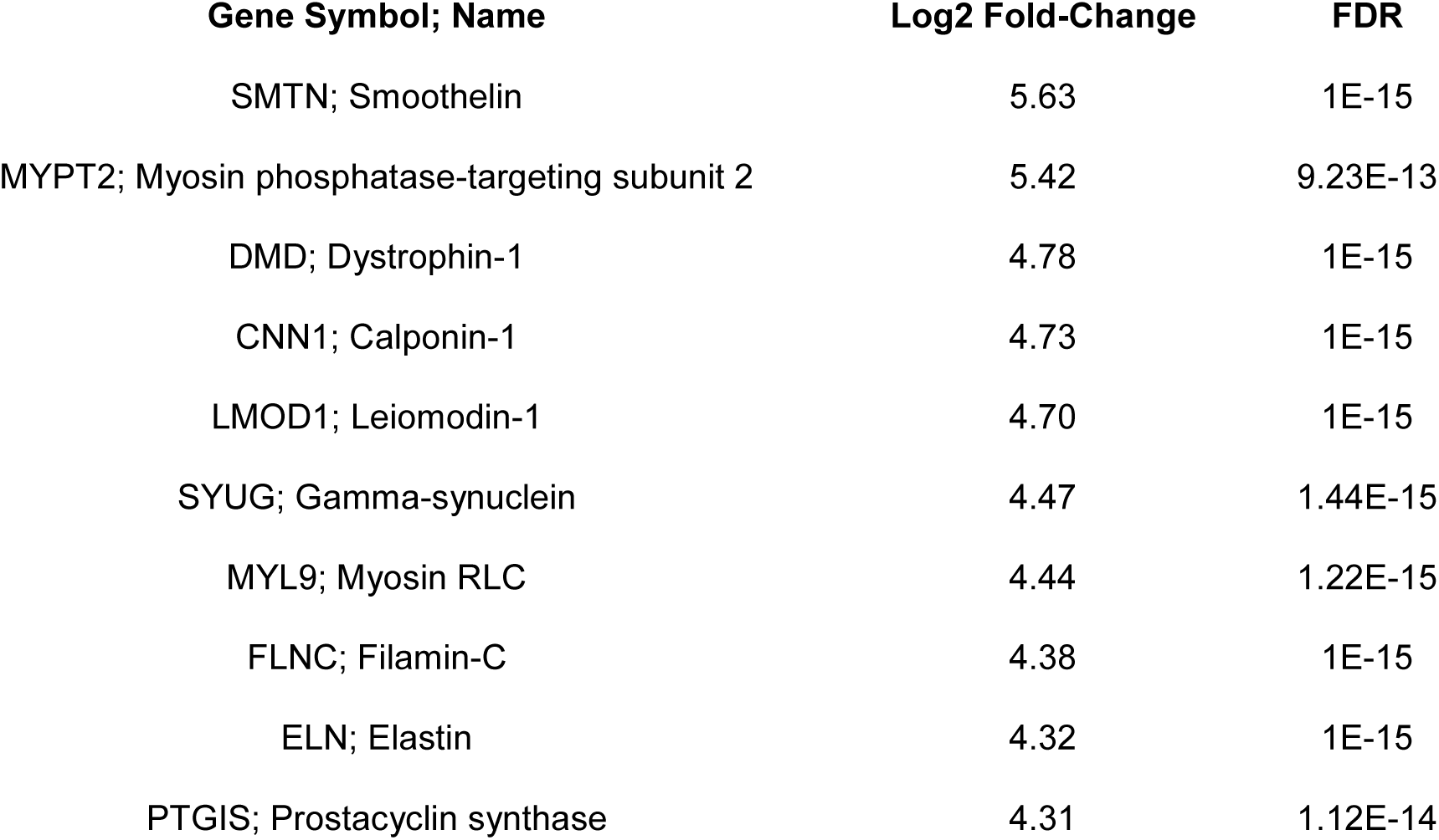
Proteins enriched in blood vessels.

**Table 3:** Reactome pathways enriched in alveoli.

| **Pathway Name/**<br>**Reactome Pathway Identifier (R-HSA)** | **Log2 Fold-**<br>**Change** | **FDR** | **Set Size** |
| --- | --- | --- | --- |
| Neutrophil degranulation/<br>R-HSA-6798695 | 1.06 | 1.83E-14 | 162 |
| Regulation of expression of SLITs and<br>ROBOs/ R-HSA-9010553 | 0.99 | 1.83E-14 | 283 |
| L13a-mediated translational silencing of<br>Ceruloplasmin expression/ R-HSA-156827 | 1.19 | 1.83E-14 | 61 |
| Translation initiation complex formation/<br>R-HSA-72649 | 1.21 | 1.83E-14 | 33 |
| Formation of a pool of free 40S subunits/<br>R-HSA-72689 | 1.20 | 1.83E-14 | 55 |
| Formation of the ternary complex, and<br>subsequently, the 43S complex/<br>R-HSA-72695 | 1.22 | 1.83E-14 | 29 |
| Ribosomal scanning and start codon<br>recognition/ R-HSA-72702 | 1.20 | 1.83E-14 | 34 |
| GTP hydrolysis and joining of the 60S<br>ribosomal subunit/R-HSA-72706 | 1.18 | 1.83E-14 | 62 |
| mRNA Splicing - Major Pathway/<br>R-HSA-72163 | 1.23 | 1.83E-14 | 42 |
| SRP-dependent cotranslational protein<br>targeting to membrane/R-HSA-1799339 | 1.23 | 1.83E-14 | 56 |

**Table 4:** Proteins over-represented in alveoli.

| **Gene Symbol; Name** | **Log2 Fold-Change** | **FDR** |
| --- | --- | --- |
| CAH4; Carbonic anhydrase | 5.11 | 1.22E-15 |
| ACSL5; Long-chain-fatty-acid CoA ligase 5 | 4.50 | 6.11E-13 |
| AMPE; Aminopeptidase A | 4.37 | 1.04E-09 |
| AQP4; Aquaporin-4 | 4.32 | 1.65E-14 |
| RAGE; Advanced glycosylation end product-specific<br>receptor | 4.24 | 7.52E-14 |
| ABCA3; ATP-binding cassette sub-family A member 3 | 4.19 | 1.65E-14 |
| K2C7; Cytokeratin-7 | 4.12 | 1.65E-14 |
| K1C18; Cytokeratin-18 | 3.90 | 1.65E-14 |
| PCAT1; LPC acyltransferase 1 | 3.80 | 1.65E-14 |
| K1C19; Cytokeratin-19 | 3.75 | 4.17E-05 |

We next specifically examined the core-matrisome (red dots) and matrisome-associated (blue dots) proteins (**Fig 3B**). Consistent with prior knowledge, blood vessels have been shown to be enriched in aggrecan (ACAN) elastin (ELN), emilin (EMIL1), lumican (LUM), tenascin (TNC), von Willebrand factor (VWF), nidogen (NID1), and collagen VIII (CO8A2) which are highlighted in the volcano plot (33). These data further support that our microproteomics protocol can delineate the ECM composition between distinct regions.

## DISCUSSION

Common proteomics-based efforts to map tissue composition are limited by the loss of spatial information caused by the need to completely homogenize tissue pieces during sample preparation. Here we developed a novel micro-proteomics strategy using tissue processed by the most widely used staining technique to identify regions of interest, followed by microdissection and subsequent proteomics. We determined the ECM composition of morphologically normal lung alveolar structures as compared to adjacent human lung blood vessels in unaffected regions from IPF lungs. To address the many challenges including low tissue volumes (<0.1 mm^3^), formalin-fixation, and histological stains, we utilized a variety of strategies employing commercially available tools to yield 1107 and 683 unique proteins to morphologically normal human lung alveoli and human lung blood vessels, respectively, and identified a total of 137 as ECM proteins. Using BayesENproteomics (23), we identify 1252 differentially expressed proteins between these regions. This is an improvement from a previous study using H&E stained FFPE tissue. In this study, researchers used 0.5 mm^3^ of tissue (5X more starting material than the current study), to identify up to 714 unique proteins (14). However, this study used a threshold of two or more unique identifying peptides to identify proteins, whereas the current study applies a more stringent threshold of three or more unique identifying peptides to identify proteins.

An emerging theme is that the ECM is a driver of disease processes including atherosclerosis (34) fibrosis (35, 36) and cancer (37). The strategy developed here could be applied to a multitude of settings where tissue heterogeneity is a common theme. Currently, ECM tissue atlases of IPF (38) and, to an extent, liver fibrosis (39) have been developed to help researchers better understand and model fibrosis progression. Thus, this strategy could be applied to archived FFPE tissues to reliably determine not only regional ECM composition, but cellular pathways perturbed in health and disease.

The work described here could be enhanced by the combination of other ‘omic’ studies. For instance, serial sections following laser capture microdissection could be used for next-generation sequencing of RNA or DNA (40) for a more complete profiling of the regions of interest. In addition, matrix-assisted laser desorption/ionization (MALDI) could be applied to determine gradient changes at defined tissue interphases (41). A limitation to this study is that H&E staining relies on pattern recognition rather than staining for specific proteins, however, our approach could be combined with spatially targeted optical microproteomics (STOMP) which combines antibodies and fluorescence to identify regions of interest (42).

Our work is a step towards interrogating cellular pathways and ECM changes within complex tissues. The application of this novel microproteomics protocol, using commercially available tools, will enhance the development of comprehensive tissue atlases for a variety of pathologies.

## ACKNOWLEDGEMENTS

The authors would like to thank Peter Walker at the University of Manchester Histology Core Facility and Roger Meadows at the University of Manchester BioImaging facility for the histological and laser capture microscopy support, respectively. In addition, we acknowledge the Wythenshawe Hospital Transplant Unit and Prof. James Fildes for helping us acquire human lung specimens for research.

## AUTHOR CONTRIBUTIONS

JAH and VM conducted all data analysis. JAH and SR performed the mass spectrometry preparations. JAH and AM performed and validated the histology. SW, RO, DK optimized the mass spectrometry. JAH, VM, MAS, JS wrote the manuscript with all author inputs. JAH conceived the project and supervised all experiments.

## DECLARATION

The authors have declared that no conflict of interests exists.

---

**Supplemental Figure 1:**
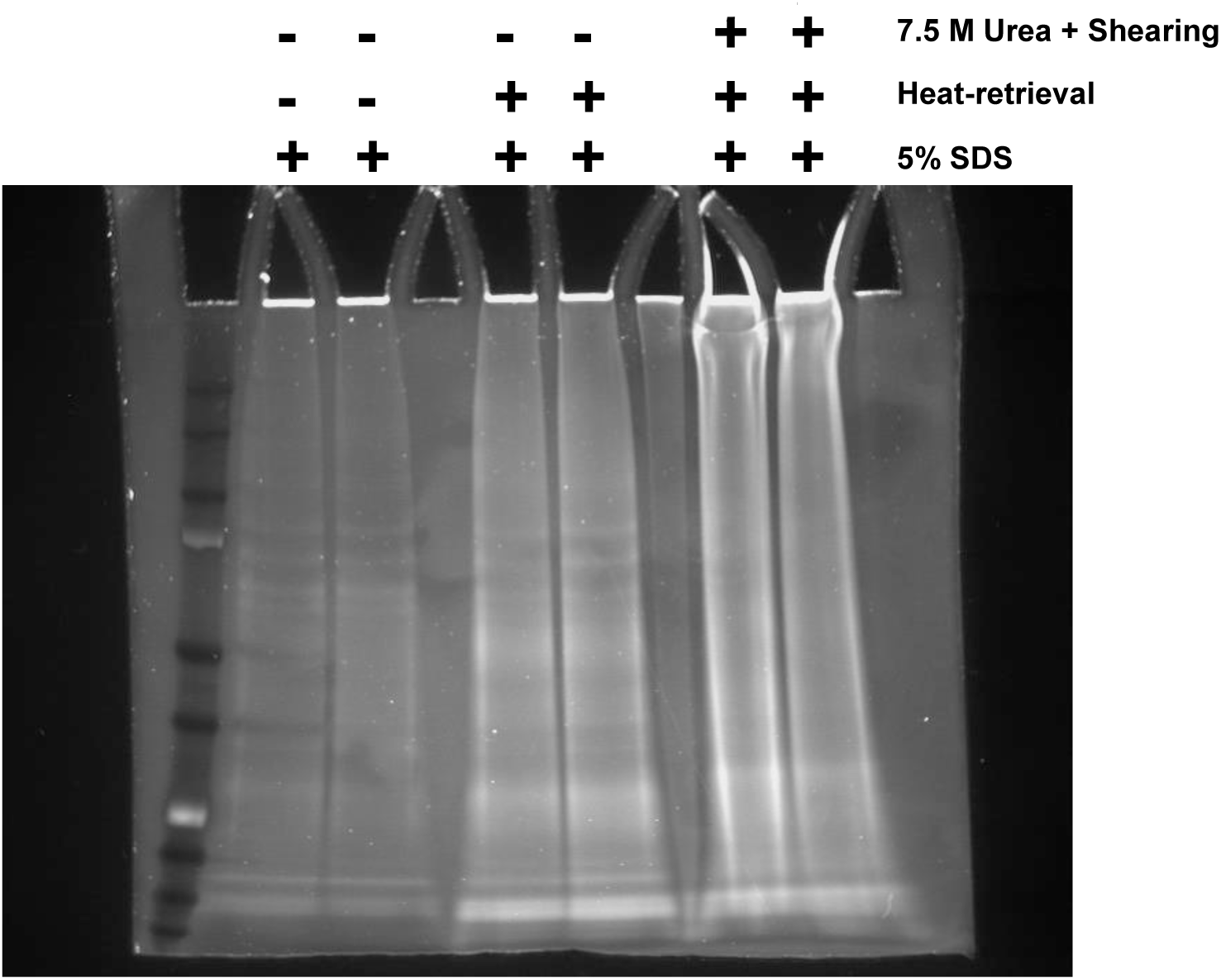
Protein extraction of H&E stained FFPE tissue sections. A 5-micron section of IPF tissue was serially sectioned and H&E stained. The whole tissue was used and subjected to 5% SDS alone, 5% SDS with heat-treatment, or 5% SDS treatment with heat-treatment followed by shearing in the presence of 7.5 M urea. Shown is a Sypro Ruby SDS-PAGE gel of complete lysates.

