## Supplemental Table 1 for "Laser capture microdissection coupled mass spectrometry (LCM-MS) for spatially resolved analysis of formalin-fixed and stained human lung tissues"

| **UniProt** | **Protein name** | **Gene symbol** | **Accession** |
| --- | --- | --- | --- |
| Q9Y6C2 | EMILIN-1 (Elastin microfibril interface-located protein 1) (Elastin microfibril interfacer 1) | *EMILIN1* | EMIL1 |
| P02452 | Collagen alpha-1(I) chain (Alpha-1 type I collagen) | *COL1A1* | CO1A1 |
| P02461 | Collagen alpha-1(III) chain | *COL3A1* | CO3A1 |
| P02462 | Collagen alpha-1(IV) chain [Cleaved into: Arresten] | *COL4A1* | CO4A1 |
| P05997 | Collagen alpha-2(V) chain | *COL5A2* | CO5A2 |
| P12110 | Collagen alpha-2(VI) chain | *COL6A2* | CO6A2 |
| P12111 | Collagen alpha-3(VI) chain | *COL6A3* | CO6A3 |
| P27658 | Collagen alpha-1(VIII) chain (Endothelial collagen) [Cleaved into: Vastatin] | *COL8A1* | CO8A1 |
| P12107 | Collagen alpha-1(XI) chain | *COL11A1* | COBA1 |
| Q99715 | Collagen alpha-1(XII) chain | *COL12A1* | COCA1 |
| P01040 | Cystatin-A (Cystatin-AS) (Stefin-A) [Cleaved into: Cystatin-A, N-terminally processed] | *CSTA* | CYTA |
| P08311 | Cathepsin G (CG) (EC 3.4.21.20) | *CTSG* | CATG |
| Q9UBR2 | Cathepsin Z (EC 3.4.18.1) (Cathepsin P) (Cathepsin X) | *CTSZ* | CATZ |
| P07585 | Decorin (Bone proteoglycan II) (PG-S2) (PG40) | *DCN* | PGS2 |
| Q8IUX7 | Adipocyte enhancer-binding protein 1 (AE-binding protein 1) (Aortic carboxypeptidase-like protein) | *AEBP1* | AEBP1 |
| P30740 | Leukocyte elastase inhibitor (LEI) (Monocyte/neutrophil elastase inhibitor) (EI) (M/NEI) (Peptidase inhibitor 2) (PI-2) | *SERPINB1* | ILEU |
| P23142 | Fibulin-1 (FIBL-1) | *FBLN1* | FBLN1 |
| P98095 | Fibulin-2 (FIBL-2) | *FBLN2* | FBLN2 |
| P35555 | Fibrillin-1 [Cleaved into: Asprosin] | *FBN1* | FBN1 |
| P02671 | Fibrinogen alpha chain [Cleaved into: Fibrinopeptide A; Fibrinogen alpha chain] | *FGA* | FIBA |
| P02675 | Fibrinogen beta chain [Cleaved into: Fibrinopeptide B; Fibrinogen beta chain] | *FGB* | FIBB |
| Q14112 | Nidogen-2 (NID-2) (Osteonidogen) | *NID2* | NID2 |
| Q63ZY3 | KN motif and ankyrin repeat domain-containing protein 2 (Ankyrin repeat domain-containing protein 25) (Matrix-remodeling-associated protein 3) | *KANK2* | KANK2 |
| P01023 | Alpha-2-macroglobulin (Alpha-2-M) (C3 and PZP-like alpha-2-macroglobulin domain-containing protein 5) | *A2M* | A2MG |
| P04083 | Annexin A1 (Annexin I) (Annexin-1) (Calpactin II) (Calpactin-2) (Chromobindin-9) (Lipocortin I) | *ANXA1* | ANXA1 |
| P12429 | Annexin A3 (35-alpha calcimedin) (Annexin III) (Annexin-3) (Inositol 1,2-cyclic phosphate 2-phosphohydrolase) (Lipocortin III) | *ANXA3* | ANXA3 |
| P09525 | Annexin A4 (35-beta calcimedin) (Annexin IV) (Annexin-4) (Carbohydrate-binding protein p33/p41) (Chromobindin-4) (Endonexin I) | *ANXA4* | ANXA4 |
| P08758 | Annexin A5 (Anchorin CII) (Annexin V) (Annexin-5) (Calphobindin I) (CBP-I) (Endonexin II) (Lipocortin V) | *ANXA5* | ANXA5 |
| P02790 | Hemopexin (Beta-1B-glycoprotein) | *HPX* | HEMO |
| P04196 | Histidine-rich glycoprotein (Histidine-proline-rich glycoprotein) (HPRG) | *HRG* | HRG |
| O00468 | Agrin [Cleaved into: Agrin N-terminal 110 kDa subunit; Agrin C-terminal 110 kDa subunit; Agrin C-terminal 90 kDa fragment (C90) | *AGRN* | AGRIN |
| Q86YZ3 | Hornerin | *HRNR* | HORN |
| Q5D862 | Filaggrin-2 (FLG-2) (Intermediate filament-associated and psoriasis-susceptibility protein) (Ifapsoriasin) | *FLG2* | FILA2 |
| O15230 | Laminin subunit alpha-5 (Laminin-10 subunit alpha) (Laminin-11 subunit alpha) (Laminin-15 subunit alpha) | *LAMA5* | LAMA5 |
| P09382 | Galectin-1 (Gal-1) (14 kDa laminin-binding protein) (HLBP14) (14 kDa lectin) (Beta-galactoside-binding lectin L-14-I) (Galaptin) | *LGALS1* | LEG1 |
| Q14767 | Latent-transforming growth factor beta-binding protein 2 (LTBP-2) | *LTBP2* | LTBP2 |
| P51884 | Lumican (Keratan sulfate proteoglycan lumican) (KSPG lumican) | *LUM* | LUM |
| P14543 | Nidogen-1 (NID-1) (Entactin) | *NID1* | NID1 |
| P20774 | Mimecan (Osteoglycin) (Osteoinductive factor) (OIF) | *OGN* | MIME |
| P01009 | Alpha-1-antitrypsin (Alpha-1 protease inhibitor) (Alpha-1-antiproteinase) (Serpin A1) | *SERPINA1* | A1AT |
| P35237 | Serpin B6 (Cytoplasmic antiproteinase) (CAP) (Peptidase inhibitor 6) (PI-6) (Placental thrombin inhibitor) | *SERPINB6* | SPB6 |
| P51888 | Prolargin (Proline-arginine-rich end leucine-rich repeat protein) | *PRELP* | PRELP |
| P26447 | Protein S100-A4 (Calvasculin) (Metastasin) (Placental calcium-binding protein) (Protein Mts1) (S100 calcium-binding protein A4) | *S100A4* | S10A4 |
| P05109 | Protein S100-A8 (Calgranulin-A) (Calprotectin L1L subunit) (Cystic fibrosis antigen) (CFAG) (Leukocyte L1 complex light chain) | *S100A8* | S10A8 |
| P06702 | Protein S100-A9 (Calgranulin-B) (Calprotectin L1H subunit) (Leukocyte L1 complex heavy chain) | *S100A9* | S10A9 |
| P29508 | Serpin B3 (Protein T4-A) (Squamous cell carcinoma antigen 1) (SCCA-1) | *SERPINB3* | SPB3 |
| P21980 | Protein-glutamine gamma-glutamyltransferase 2 (EC 2.3.2.13) (Tissue transglutaminase) (Transglutaminase C) (TG(C)) (TGC) | *TGM2* | TGM2 |
| Q08188 | Protein-glutamine gamma-glutamyltransferase E (EC 2.3.2.13) (Transglutaminase E) (TG(E)) (TGE) (TGase E) (Transglutaminase-3) | *TGM3* | TGM3 |
| Q05707 | Collagen alpha-1(XIV) chain (Undulin) | *COL14A1* | COEA1 |
| P04004 | Vitronectin (VN) (S-protein) (Serum-spreading factor) (V75) [Cleaved into: Vitronectin V65 subunit; Vitronectin V10 subunit] | *VTN* | VTNC |
| P04275 | von Willebrand factor (vWF) [Cleaved into: von Willebrand antigen 2 (von Willebrand antigen II)] | *VWF* | VWF |
| Q96P63 | Serpin B12 | *SERPINB12* | SPB12 |
| P55083 | Microfibril-associated glycoprotein 4 | *MFAP4* | MFAP4 |
| P49257 | Protein ERGIC-53 (ER-Golgi intermediate compartment 53 kDa protein) (Gp58) (Intracellular mannose-specific lectin MR60) | *LMAN1* | LMAN1 |
| P55268 | Laminin subunit beta-2 (Laminin B1s chain) (Laminin-11 subunit beta) (Laminin-14 subunit beta) (Laminin-15 subunit beta) | *LAMB2* | LAMB2 |
| P02751 | Fibronectin (FN) (Cold-insoluble globulin) (CIG) [Cleaved into: Anastellin; Ugl-Y1; Ugl-Y2; Ugl-Y3] | *FN1* | FINC |
| P50454 | Serpin H1 (47 kDa heat shock protein) (Arsenic-transactivated protein 3) (AsTP3) (Cell proliferation-inducing gene 14 protein) | *SERPINH1* | SERPH |
| P17931 | Galectin-3 (Gal-3) (35 kDa lectin) (Carbohydrate-binding protein 35) (CBP 35) (Galactose-specific lectin 3) | *LGALS3* | LEG3 |
| Q9UBX5 | Fibulin-5 (FIBL-5) (Developmental arteries and neural crest EGF-like protein) (Dance) (Urine p50 protein) (UP50) | *FBLN5* | FBLN5 |
| P01011 | Alpha-1-antichymotrypsin (ACT) (Cell growth-inhibiting gene 24/25 protein) (Serpin A3) | *SERPINA3* | AACT |
| P20908 | Collagen alpha-1(V) chain | *COL5A1* | CO5A1 |
| P01008 | Antithrombin-III (ATIII) (Serpin C1) | *SERPINC1* | ANT3 |
| P11047 | Laminin subunit gamma-1 (Laminin B2 chain) (Laminin-1 subunit gamma) (Laminin-10 subunit gamma) (Laminin-11 subunit gamma) | *LAMC1* | LAMC1 |
| P98160 | Basement membrane-specific heparan sulfate proteoglycan core protein (HSPG) (Perlecan) (PLC) | *HSPG2* | PGBM |
| P13611 | Versican core protein (Chondroitin sulfate proteoglycan core protein 2) (Chondroitin sulfate proteoglycan 2) | *VCAN* | CSPG2 |
| P08572 | Collagen alpha-2(IV) chain [Cleaved into: Canstatin] | *COL4A2* | CO4A2 |
| P08123 | Collagen alpha-2(I) chain (Alpha-2 type I collagen) | *COL1A2* | CO1A2 |
| P12109 | Collagen alpha-1(VI) chain | *COL6A1* | CO6A1 |
| P08133 | Annexin A6 (67 kDa calelectrin) (Annexin VI) (Annexin-6) (Calphobindin-II) (CPB-II) (Chromobindin-20) (Lipocortin VI) | *ANXA6* | ANXA6 |
| P07339 | Cathepsin D (EC 3.4.23.5) [Cleaved into: Cathepsin D light chain; Cathepsin D heavy chain] | *CTSD* | CATD |
| Q08380 | Galectin-3-binding protein (Basement membrane autoantigen p105) (Lectin galactoside-binding soluble 3-binding protein) | *LGALS3BP* | LG3BP |
| Q15582 | Transforming growth factor-beta-induced protein ig-h3 (Beta ig-h3) (Kerato-epithelin) | *TGFBI* | BGH3 |
| P02679 | Fibrinogen gamma chain | *FGG* | FIBG |
| P36955 | Pigment epithelium-derived factor (PEDF) (Cell proliferation-inducing gene 35 protein) (EPC-1) (Serpin F1) | *SERPINF1* | PEDF |
| P19823 | Inter-alpha-trypsin inhibitor heavy chain H2 (ITI heavy chain H2) (ITI-HC2) (Inter-alpha-inhibitor heavy chain 2) | *ITIH2* | ITIH2 |
| P15502 | Elastin (Tropoelastin) | *ELN* | ELN |
| P00488 | Coagulation factor XIII A chain (Coagulation factor XIIIa) (EC 2.3.2.13) (Protein-glutamine gamma-glutamyltransferase A chain) | *F13A1* | F13A |
| P20073 | Annexin A7 (Annexin VII) (Annexin-7) (Synexin) | *ANXA7* | ANXA7 |
| P21810 | Biglycan (Bone/cartilage proteoglycan I) (PG-S1) | *BGN* | PGS1 |
| P00734 | Prothrombin (EC 3.4.21.5) (Coagulation factor II) | *F2* | THRB |
| P19827 | Inter-alpha-trypsin inhibitor heavy chain H1 (ITI heavy chain H1) (ITI-HC1) (Inter-alpha-inhibitor heavy chain 1) | *ITIH1* | ITIH1 |
| Q9NVD7 | Alpha-parvin (Actopaxin) (CH-ILKBP) (Calponin-like integrin-linked kinase-binding protein) | *PARVA* | PARVA |
| Q15063 | Periostin (PN) (Osteoblast-specific factor 2) (OSF-2) | *POSTN* | POSTN |
| P39060 | Collagen alpha-1(XVIII) chain [Cleaved into: Endostatin; Non-collagenous domain 1 (NC1)] | *COL18A1* | COIA1 |
| P60903 | Protein S100-A10 (Calpactin I light chain) (Calpactin-1 light chain) (Cellular ligand of annexin II) | *S100A10* | S10AA |
| P50995 | Annexin A11 (56 kDa autoantigen) (Annexin XI) (Annexin-11) (Calcyclin-associated annexin 50) (CAP-50) | *ANXA11* | ANX11 |
| P06703 | Protein S100-A6 (Calcyclin) (Growth factor-inducible protein 2A9) (MLN 4) (Prolactin receptor-associated protein) (PRA) | *S100A6* | S10A6 |
| P07355 | Annexin A2 (Annexin II) (Annexin-2) (Calpactin I heavy chain) (Calpactin-1 heavy chain) (Chromobindin-8) (Lipocortin II) | *ANXA2* | ANXA2 |
